# Effects of Repeat Test Exposure on Gait Parameters in Naïve Lewis Rats

**DOI:** 10.1101/2023.04.19.537488

**Authors:** Nat A. Thurlow, Kiara M. Chan, Taylor D. Yeater, Kyle D. Allen

**Author notes:** Corresponding Author: Kyle D. Allen Associate Professor, J. Crayton Pruitt Family Department of Biomedical Engineering, University of Florida 1275 Center Drive, Biomedical Sciences Building Gainesville, Florida 32611. **Authors’ Contributions:** KMC, TDY, and KDA conceptualized and designed the experiment. KMC and TDY were responsible for rodent care and gait data collection. Data was processed and analyzed by NAT with input from KMC and KDA. The final transcript was written by NAT with input and approval from all authors.

## Abstract

Rodent gait analysis has emerged as a powerful, quantitative behavioral assay to characterize the pain and disability associated with movement-related disorders. In other behavioral assays, the importance of acclimation and the effect of repeated testing have been evaluated. However, for rodent gait analysis, the effects of repeated gait testing and other environmental factors have not been thoroughly characterized. In this study, fifty-two naïve male Lewis rats ages 8 to 42 weeks completed gait testing at semi-random intervals for 31 weeks. Gait videos and force plate data were collected and processed using a custom MATLAB suite to calculate velocity, stride length, step width, percentage stance time (duty factor), and peak vertical force data. Exposure was quantified as the number of gait testing sessions. Linear mixed effects models were used to evaluate the effects of velocity, exposure, age, and weight on animal gait patterns. Relative to age and weight, repeated exposure was the dominant parameter affecting gait variables with significant effects on walking velocity, stride length, fore and hind limb step width, fore limb duty factor, and peak vertical force. From exposure 1 to 7, average velocity increased by approximately 15 cm/s. Together, these data indicate arena exposure had large effects on gait parameters and should be considered in acclimation protocols, experimental design, and subsequent data analysis of rodent gait data.

## 1. Introduction

Rodent gait analysis is a powerful preclinical assay to characterize behavioral changes associated with movement-related pain and disability. Gait analysis for rodent models has been particularly useful for identifying behavioral changes that manifest due to Parkinson’s disease (Amende et al., 2005; Boix et al., 2018), spinal cord injury (Isvoranu et al., 2021; Pukale et al., 2021), osteoarthritis (Chan et al., 2022; Jacobs et al., 2014; Lakes and Allen, 2018), and other neurodevelopmental disorders (Akula et al., 2020). Detecting gait changes in rodents is not trivial, as gait analysis must be highly sensitive due to the small size and rapidity of the rodent gait sequence in comparison to humans. Moreover, rodents tend to consciously mask injuries and pain-related behaviors to avoid attention from predators (Lakes and Allen, 2016; Sheppard et al., 2022; Stasiak et al., 2003). Advanced motion tracking technology, ground reaction force measurements, and computational tools have accelerated the application of gait analysis for quantitative, objective measurements in rodent models; however, analyzing rodent gait data remains complex and nuanced (Batka et al., 2014; Jacobs et al., 2018; Jacobs and Allen, 2020).

Critical sources of variation, such as walking velocity, animal sex, strain, housing environment, body weight, and age, need to be controlled during experimental design and statistical analysis. For example, the relationship between velocity and gait have been well-established, with higher velocities causing longer stride lengths and shorter stance times (Batka et al., 2014; Górska et al., 1998; Westerga and Gramsbergen, 1993). Unfortunately, even velocity effects are not always addressed during the analysis of rodent gait (Lakes and Allen, 2016). Like velocity, body weight and animal age can also affect gait measures (Akula et al., 2020; Bair et al., 2019; Jacobs and Allen, 2020; Pitzer et al., 2021). Here, body weight is often used as a correlate to an animal’s skeletal size, where weight can affect spatial gait parameters such as stride length and step width (Pitzer et al., 2021; Sheppard et al., 2022). Similarly, age affects both velocity over time and disease-related shifts in gait parameters (Bair et al., 2019; Broom et al., 2021; Chan et al., 2022). Careful consideration of these factors through experimental design and data analysis is key to interpreting gait data.

While velocity, weight, and age have known effects on gait, these factors are also interrelated, as weight increases with age and velocity fluctuates throughout life. Moreover, there are other relevant, related factors beyond velocity, weight, and age that are not commonly assessed. The effects of test conditioning have been established for other behavioral assays, but there is far less literature and discussion on repeated testing and the role of arena exposure in rodent gait analysis. The objective of this study is to analyze the effects of repeated testing and arena exposure on spatial, temporal, and dynamic gait parameters relative to the effects of age and weight. In doing so, we highlight the need to consider repeated testing exposure when assessing rodent gait and suggest ways to include exposure in data visualization and analysis.

## 2. Methods

### 2.1 Animals

In this study, 52 naïve male Lewis rats were acquired between ages 8 to 42 weeks and underwent gait testing at semi-random intervals over a 31-week period. The testing schedule varied for each animal, with the gait testing frequency ranging from twice per week to once every four weeks. In total, each animal was tested at least 6 times. The original experimental design was set to provide extended acclimation for a separate age-related experiment and create a large database of age-weight matched control data (Chan et al., 2022). Age and weight were recorded at the time of testing. Exposure was quantified from the testing schedule where one arena exposure was defined as a single gait testing session. Details of the animals’ age and weight at their first and last exposure are included in Supplementary Table 1. Animals were housed in pairs with ad libitum access to food and water in an atmosphere-controlled room with a 12h light dark cycle. All methods followed the recommendations set by the Association for Assessment and Accreditation of Laboratory Animal Care and were approved by the University of Florida Institutional Animal Care and Use Committee.

### 2.2 Gait Testing

Features of the gait arena, configuration of the video recording system, and the MATLAB suite used to process video and force plate data are extensively detailed in previous work (Jacobs et al., 2018; Kloefkorn et al., 2017). Briefly, the gait arena consists of a 60×5×10 inch acrylic enclosure (gait arena) mounted on an aluminum frame. Inside the aluminum base, a mirror angled at 45 degrees and positioned beneath the arena floor enables visualization of both the lateral and ventral views of an animal. Four regions of the arena were instrumented with force recording equipment as described in Jacobs, et al. To collect gait data, animals were individually placed in the gait arena and allowed to freely explore at self-selected walking speeds. Each trial consists of simultaneous high-speed video (500 frames/s, Phantom Miro Lab 320) and force plate signal collection (Kistler 3-Component Force Link Type 9317B) as the rat travels across the arena at a constant, self-selected velocity. Each gait testing session (defined as one exposure) lasted 20-30 minutes per animal, with four to sixteen trials collected per animal per session. Videos and force data were processed using Automated Gait Analysis through Hues and Areas version 2 (AGATHAv2) (Kloefkorn et al., 2017) and dynamicEDGAR pipelines for calculating spatiotemporal and dynamic parameters (Jacobs et al., 2014). For a list and explanation of these parameters see Jacobs, et al., 2014. Age, weight, and the number of arena exposures at testing were included as identifiers with these data.

### 2.3 Statistical Analysis

For consistent visualization and statistical analysis, age, weight, and exposure number were converted into discrete classes and treated as grouping factors. Age groups were separated into roughly 7-week increments containing a similar number of total trials. The age classes were 8-14 weeks, 15-21 weeks, 22-28 weeks, 29-35 weeks, and 36-57 weeks. The weight classes were similarly divided to consist of less than 350 grams, 350-400 grams, 400-450 grams, 450-500 grams, and greater than 500 grams. Exposures 1 through 7 were included in the final data analysis as there were an insufficient number of trials collected for exposure 8 and beyond. A total of 694 trials were analyzed for spatiotemporal gait parameters and 791 force data curves were analyzed for dynamic gait parameters.

Data visualization and statistical analysis were conducted in R (version 4.0.2). For stride length, duty factor, step width, and peak vertical force, linear mixed effects models were used to simultaneously assess the fixed effects of velocity, weight, age, and exposure. For velocity, a linear mixed effects model was used to simultaneously assess the fixed effect of weight, age, and exposure. In all linear mixed effects models, animal identifiers were used to capture the random effect of repeated testing on the same animal. Models did not include interactions among the fixed effects, as goodness of fit (R^2^) was collected for each model. The significance of fixed effects was assessed using a type III analysis of variance using Satterthwaite’s method. When indicated by the analysis of variance, comparisons of least square means estimates within a given fixed effect were conducted using a pairwise, Tukey’s HSD test for multiple comparisons. In all statistical tests, a p-value of 0.05 or lower was considered statistically significant. Throughout the paper, data are presented as mean ± 95% confidence intervals.

## 3. Results

### 3.1 Velocity

Animals walked at faster self-selected velocities as the number of arena exposures increased (p<0.001) and slowed slightly as age increased (p=0.0011, Fig. 1). The relationship between exposure and velocity was linear with an incremental increase following each exposure (Figure 1). In total, average velocity increased by 38.1% from 37.87 ± 2.33 cm/s to 52.31 ± 2.58 cm/s after 7 exposures to the gait testing arena (Fig. 1A). At exposure 1 and exposure 2, animals walked at significantly lower velocities than all subsequent exposures (p<0.05). Velocity at exposure 3 was also lower than exposures 5 through 7 (p<0.01). Although velocity continually increased, the mean differences across exposures 4 through 7 were less pronounced.

**Fig. 1.**
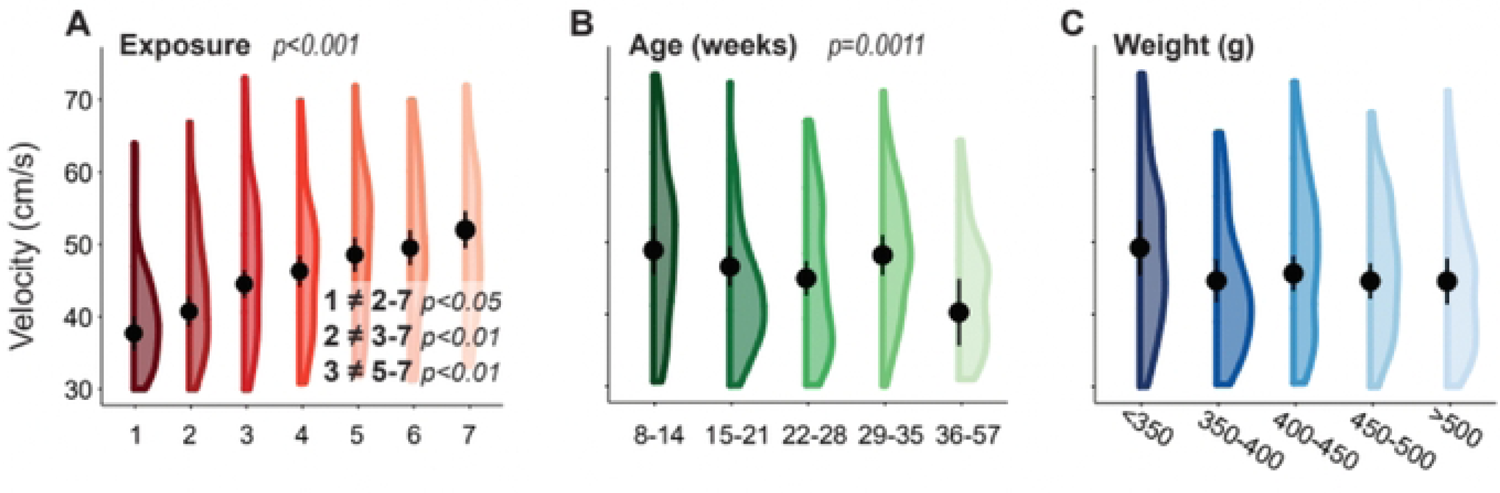
Each violin plot reports the means, 95% confidence intervals, and distributions of self-selected velocities based on linear mixed effects models for exposure (A), age (B), and weight (C). Significant main effects and pairwise comparisions have p-values listed.

The relationship between velocity and age was less uniform but still demonstrated a slight decrease with each age group (p=0.0011, Fig. 1B). Here, mean velocity decreased by 8.68 ± 6.65 cm/s from the youngest to oldest age group (p<0.05).

Velocity slightly decreased as weight increased; however, these effects were not statistically significant (Fig. 1C).

### 3.2 Stride Length

Stride lengths generally increased with exposure (p=0.0506) and age (p<0.05, Fig. 2). There was a signficant increase from exposure 1 to exposures 2 through 7 (p<0.05, Fig. 2A). Importantly, these stride length changes stabilized after 2 exposures (exposures 3-7); however, the velocity-stride length relationships were shifted up by 0.38 ± 0.25 cm from exposure 1 to exposure 6 (p<0.05) and 0.24 ± 0.24 cm from exposure 1 to 7 (p<0.05).

**Fig. 2.**
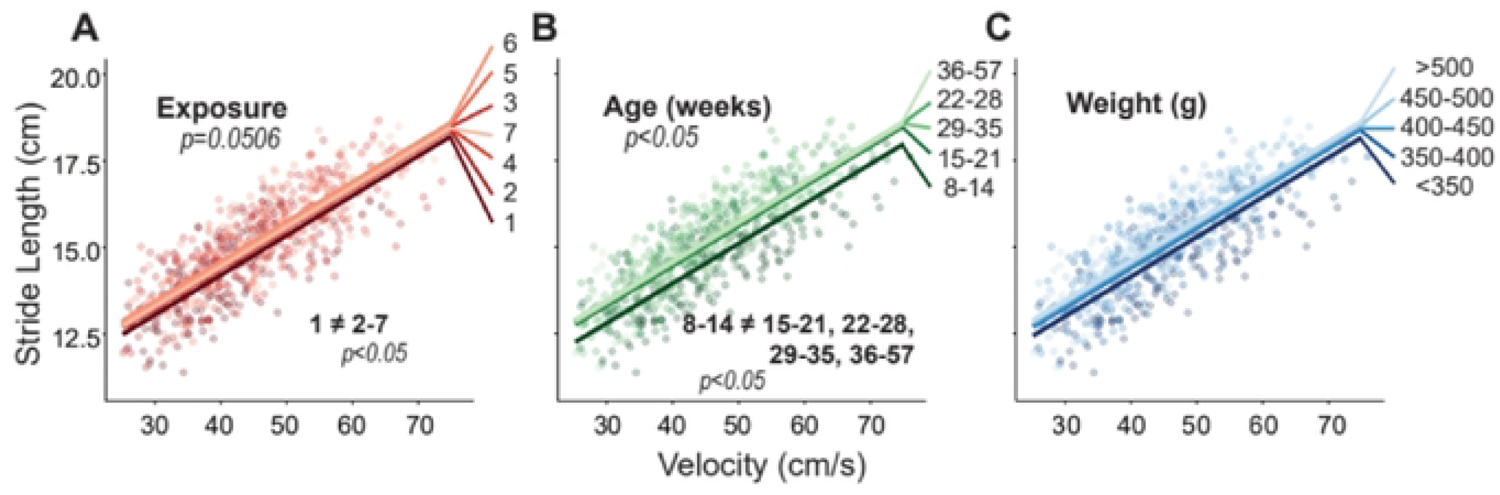
Results for stride length plotted against velocity, where raw data points indicate individual gait trials collected from each animal over the course of the entire study. Linear fits for each group are based on the linear mixed effects model for exposure (A), age (B), and weight (C). Significant main effects and pairwise comparisions have p-values listed.

Stride length became larger at older ages, with a marked difference between the 8-14 weeks group and all older groups (p<0.05, Fig. 2B). The largest difference existed between the 8-14 week and 36-57 week with a 0.63 ± 0.54 cm increase.

There was no significant effect of weight on stride length, but generally, stride length increased as weight increased (Fig. 2C).

### 3.3 Step Width

For hind limb step width, there was an effect of age (p<0.001) and weight (p<0.05) and tended to be an effect of exposure (p=0.0533); for fore limb step width, there was a signficant effect of exposure (p<0.05) and age (p<0.001, Fig. 3). Within fore step width, exposure 1 through 3 showed narrower step widths than exposure 6 (p<0.05), and exposure 2 and 3 showed narrower step width than exposure 7 (p<0.05, Fig. 3A). There were no distinct trends with exposure for hide step width.

**Fig. 3.**
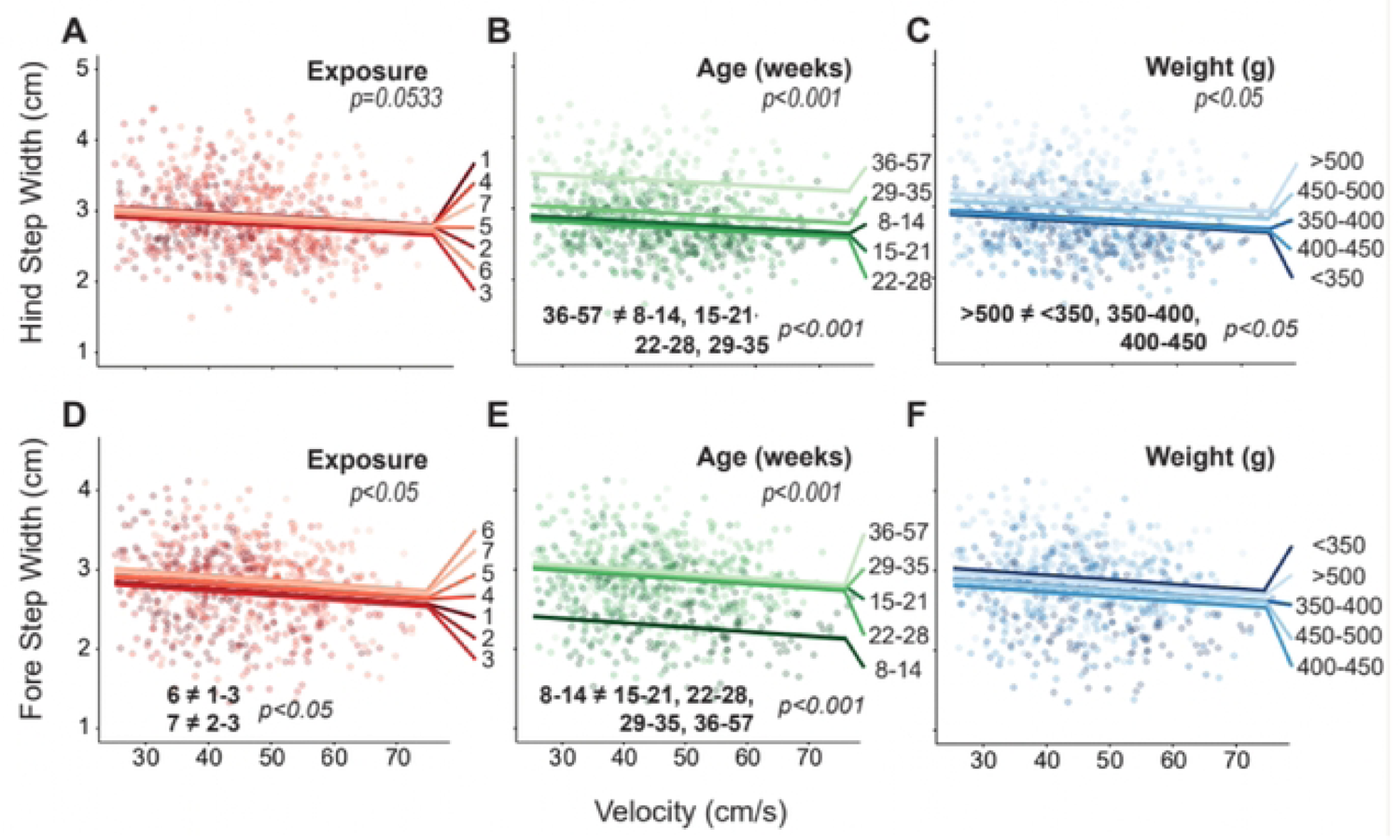
Results for hind and fore limb step width plotted against velocity, where raw data points indicate individual gait trials collected from each animal over the course of the entire study. Linear fits for each group are based on the linear mixed effects model for exposure (A, D), age (B, E), and weight (C, F). Significant main effects and pairwise comparisions have p-values listed.

For age, hind limb step width was larger in the older 36-57 week group compared to all younger groups (p<0.001) with a 0.66 ± 0.23 cm difference between the 36-57 week and 22-28 week age groups (Fig. 3B). Similarly, fore step width was larger in all older groups compared to the 8-14 week group (p<0.001) with a 0.60 ± 0.30 cm increase from the 8-14 week to the 36-57 week age group (Fig. 3E).

Hind step width also increased with weight, where the >500g weight group had wider steps than the three lightest weight groups (p<0.05, Fig. 3C). The largest difference existed between the <350G and >500g with a 0.29 ± 0.25 cm increase.

### 3.4 Duty Factor

Duty factor is defined as the amount of time the limb is in ground contact relative to total stride time. Here, duty factors are known to vary between the fore and hind limbs of most quadrupeds but are similar for the left and right limb of a given limb pair for a healthy animal. Hind limb duty factor was affected by weight (p<0.001); fore limb duty factor was affected by exposure (p<0.05), age (p<0.001), and weight (p<0.01, Fig. 4). However, despite a significant main effect, clear differences between specific numbers of exposure were not found in post-hoc tests (Fig. 4A, 4D).

**Fig. 4.**
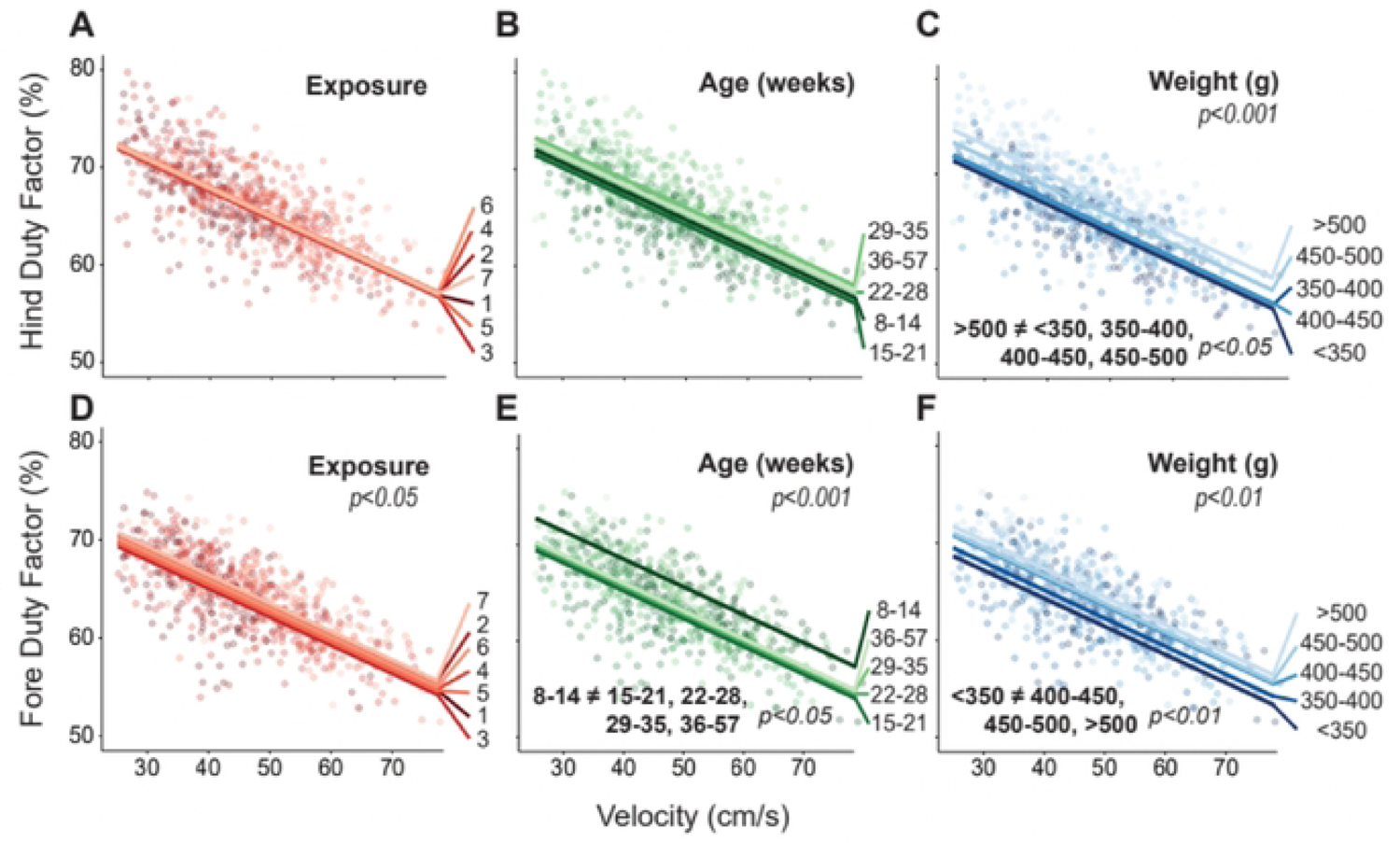
Results for hind and fore limb duty factor plotted against velocity, where raw data points indicate individual gait trials collected from each animal over the course of the entire study. Linear fits for each group are based on the linear mixed effects model for exposure (A, D), age (B, E), and weight (C, F). Significant main effects and pairwise comparisions have p-values listed.

For age, fore limb duty factors were higher in the 8-14 weeks group compared to all older groups (p<0.05) (Fig. 4E). From the 8-14 week to 36-57 week group, there was a 2.46 ± 1.86% decrease in fore limb duty factor.

For weight, hind limb duty factors were greater in the >500g group compared to all other groups accounting for a 3.17 ± 1.71% increase from the <350G to >500g group (p<0.05, Fig. 4F). Similarly, fore limb duty factors were highest in the three heaviest weight groups compared to the <350g group with a 3.06 ± 1.70% increase from the <350G to the >500g group (p<0.01, Fig. 4C).

### 3.5 Peak Vertical Force

Peak vertical force is typically plotted as a percentage of body weight. This weight-normalized peak vertical force increased with exposure (p<0.01, Fig. 5). Despite the significance of exposure, the trend between exposure number and peak vertical force is less consistent than other gait parameters, but generally showed increases from earlier to later exposures (Fig. 5A). Here, exposures 2 and 7 had the lowest velocity-peak vertical force relationship while exposures 4 through 6 had the highest (p<0.05).

**Fig. 5.**
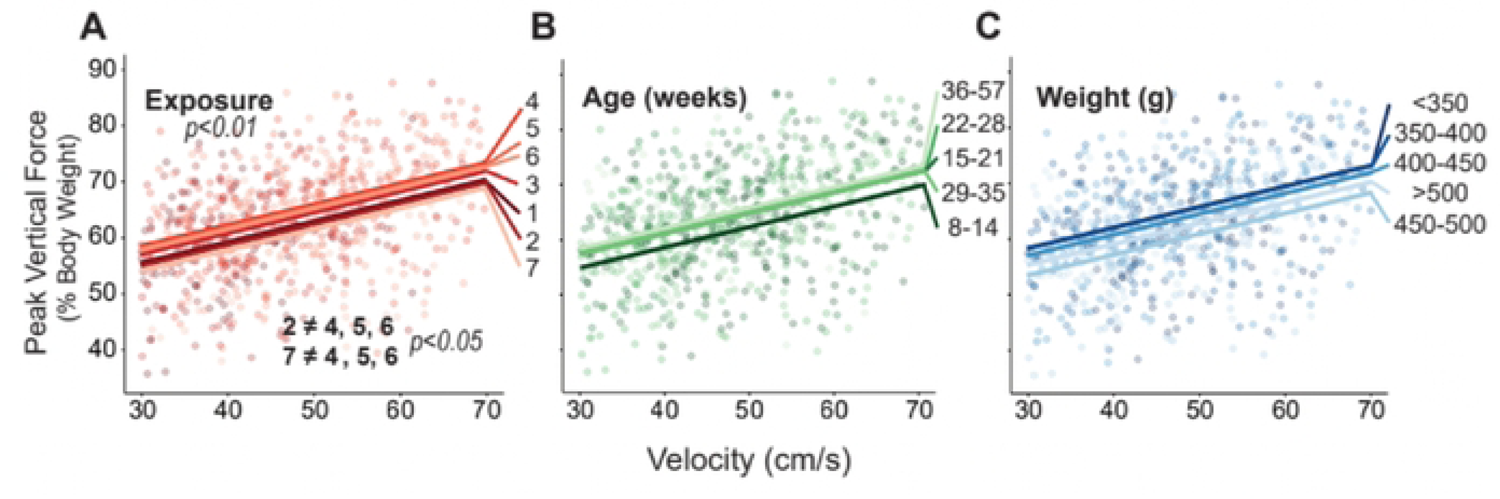
Results for peak vertical force plotted against velocity, where raw data points indicate individual gait trials collected from each animal over the course of the entire study. Linear fits for each group are based on the linear mixed effects model for exposure (A), age (B), and weight (C). Significant main effects and pairwise comparisions have p-values listed.

Although age did not have a substantial effect on peak vertical force, younger animals tended to exert less force than older age groups (Fig. 5B).

On the other hand, despite peak vertical force being normalized to weight, the heavier weight groups tended to produce lower weight-normalized peak vertical forces than the lighter weight groups (Fig. 5C); however. this fixed effect of weight was not significant in our model.

## 4. Discussion

The first objective of this study was to assess the effects of repeated arena exposures on several gait parameters. Most markedly, velocity substantially increased with repeated exposures, leading to an overall increase of 14.43 ± 3.25 cm/s after 7 exposures. Additionally, stride length increased across 7 exposures with a considerable increase from exposure 1 to all subsequent exposures. Similarly, peak vertical force generally increased across all exposures with a sizable 3.47 ± 2.53% increase from exposure 2 to 4. For step width and duty factor, there was no clear across exposure, but exposure remained a significant predictor for fore limb step width and duty factor. Considering these effects together, exposure plays a considerable role in several gait parameters with marked increases after repeated testing for velocity, stride length, and peak vertical force.

The second objective was to assess the effects of age and weight in naïve animals and compare the magnitude of exposure effects to those of age and weight. Age and weight have been analyzed as factors in previous gait work but have not been fully characterized in a large population of naïve rats (Jacobs et al., 2014; Jacobs and Allen, 2020; Pitzer et al., 2021). The effects of age were substantial for stride length, step widths, and fore duty factor. Instead of incremental increases, most significant differences related to age were between the youngest cohort and all other cohorts. For example, stride length and fore limb step width were significantly smaller in the 8-14 weeks age group compared to all older groups. Stride length, which typically ranges from 12-20cm, was about 0.55cm shorter; fore limb step width, which usually ranges from 1.5-4cm, was about 0.5cm narrower. These spatial gait changes may be attributed to the logarithmic relationship between rat size and age in which skeletal growth increases quickly from 8 to 21 weeks and plateaus following this period (Clemens et al., 2014; Hughes and Tanner, 1970; Sengupta, 2013). This likely causes spatial variables in skeletally immature rats to be distinctly smaller than their older counterparts. After this growth period, age effects may reflect degenerative aging rather than skeletal growth.

Weight drove changes in fore and hind limb duty factors and hind limb step width that reflected increased weight bearing. Hind step width and both duty factors were larger in the heaviest animals compared to all lighter groups. Compared to the three lightest groups, hind step width which typically ranges from 2-4.5cm, increased by about 2.5cm in the >500g group, and hind duty factor, which typically ranges from 55-80%, increased by about 4% in >500g group. Increased hind step width reflects a larger base of support, which may be needed to brace additional mass in the hind region of heavier rats. Higher duty factors in both limbs indicate that heavier rats spend more time in stance and thus must have faster swing times.

The effects of exposure, however, were comparable or larger than age and weight in velocity, stride length, and peak vertical force. The nearly 15cm/s increase in velocity after 7 exposures is about twice the size of the velocity decrease across age groups and three times the size of the decrease across weight groups, indicating exposure is the strongest factor for velocity. For stride length and peak vertical force, exposure, age, and weight had approximately equal shifts indicating exposure was comparable to age and weight for these parameters. As mentioned previously, the original purpose of this data collection was to establish a large age- and weight-matched database for future studies. We expected the effects of weight and age to be dominant; however, our data suggest the magnitude of exposure effects is just as critical as age and weight.

The effects of repeated test exposure have been demonstrated in other gait and behavioral testing, but the reason rodents alter their gait after repeated tests remains uncertain. In previous gait studies conducted by our group, velocity increased across testing time points regardless of surgical grouping, potentially due to repeated exposure to gait testing (Chan et al., 2022; Kloefkorn et al., 2015). Analysis of repeated test exposures has also been performed in other classical behavioral assays such as the elevated plus maze, elevated zero maze, forced swim, open-field, and light/dark exploration test, but with largely mixed results depending on the type of tests, type of animal used, and frequency of exposures (Blokland et al., 2012; Bodden et al., 2018; Tucker and McCabe, 2017). The behavioral mechanisms underlying repeat exposures have been studied extensively in these traditional rodent behavioral assays by introducing rodents to a repeated battery of tests and comparing their performance to rodents with no previous test experience. There are several proposed associations between repeated testing and altered behaviors for rodent behavioral assays including the modulation of motivation, exploratory behavior, learning, and sensitivity to stimuli (McIlwain et al., 2001; Võikar et al., 2004). Because gait testing similarly assesses behavior in rodents, it is likely that several of these behavioral links also cause changed movement patterns with repeated testing. However, gait testing is complex, and several factors may change from test to test, making the exact reasons why rodents alter their gait uncertain.

To control for size effects related to age and weight, historical data on age- and weight-matched controls or preliminary baseline gait data have been used; however, when considering the effects of repeated exposures, historical naïve data may fail to be an adequate control unless the exposure number was also explicitly recorded. Similarly, comparing post-operative or post-treatment gait data to a baseline dataset presents issues with comparing rodents at different stages of repeated testing. Further measures are needed to account for exposure effects. One method of control is rodent acclimation. Most parameters were highly affected between the first few testing exposures and showed slowed or minimal change after the 3^rd^ testing session. By completing three testing sessions before experimental data is collected, the experimenter can further minimize the effects of exposure. Internal naïve/sham controls also reduce the effects of exposure within one timepoint. By keeping gait testing exposure at each timepoint consistent between control and experimental groups, experimenters can compare groups at one timepoint. Data analysis techniques can also be used to account for increasing exposure throughout a study. Unfortunately, even if exposure is kept uniform throughout a study and across all animals, it may correlate with time and thus conclusions related to injury or disease progression. To best control for exposure effects, gait experiments should consider (1) adequately acclimating animals to the gait testing environment, (2) the inclusion of internal naïve or sham controls that complete gait testing in parallel to the experimental group, and/or (3) including testing exposure as a factor in statistical analysis.

Though we establish testing exposure as a key variable in gait analysis, considerations should be taken when applying these findings to future gait experiments. Due the original objective of this data collection, we did not anticipate the dominant effect of exposure when developing the semi-random test schedule, thus only seven testing exposures were carried out among a majority of animals. Because of this, data analysis included only up to seven exposures to ensure roughly equal trial distribution within each exposure, age, and weight grouping. However, this limits the ability to draw conclusions about long-term effects of repeated testing beyond seven sessions. Note, the velocity changes with exposure did not stabilize, even after seven exposures. Although the magnitude and significance of change for most gait parameters declined substantially by exposure 4, 5, 6 and 7, a definitive asymptote may not be reached for all gait parameters. Additionally, age, weight, and exposure are inevitably autocorrelated in the model. However, as noted in Supplemental Table 1, the age of first exposure ranged from 12 to 52 weeks old and weight at first exposure ranged from 346-662g. Despite this large range of starting ages and weights, exposure remained a dominant effect. We also selected type III analysis of variance models due to this autocorrelation, rather than a sequential type I analysis; the robustness of the model was also confirmed by switching the order the factors that appear in the model and confirming that the statistical outcomes did not change.

In conclusion, these findings establish repeated testing exposure as a substantial factor affecting gait testing in rodents. With effects comparable or greater than the effects of age and weight, it is recommended that researchers design experiments, acclimation procedures, and data analysis with the effects of repeated exposure in mind.

## Acknowledgements

This work was supported by grants from the National Institute of Health R01AR07143 and R01AR06842.

